# CD4^+^ but not CD8^+^ T cells are required for protection against severe guinea pig cytomegalovirus infections

**DOI:** 10.1101/2024.08.21.608907

**Authors:** Tyler B. Rollman, Zachary W. Berkebile, Dustin M. Hicks, Jason S. Hatfield, Priyanka Chauhan, Marco Pravetoni, Mark R. Schleiss, Gregg N. Milligan, Terry K. Morgan, Craig J. Bierle

## Abstract

Human cytomegalovirus (HCMV) is a ubiquitous herpesvirus and the leading cause of infectious disease related birth defects worldwide. How the immune response modulates the risk of intrauterine transmission of HCMV after maternal infection remains poorly understood. Maternal T cells likely play a critical role in preventing infection at the maternal-fetal interface and limiting spread across the placenta, but concerns exist that immune responses to infection may also cause placental dysfunction and adverse pregnancy outcomes. This study investigated the role of CD4^+^ and CD8^+^ T cells in a guinea pig model of primary cytomegalovirus infection. Monoclonal antibodies specific to guinea pig CD4 and CD8 were used to deplete T cells in non-pregnant and in pregnant guinea pigs after mid-gestation. CD4^+^ T cell depletion increased the severity of illness, caused significantly elevated viral loads, and increased the rate of congenital guinea pig cytomegalovirus (GPCMV) infection relative to animals treated with control antibody. CD8^+^ T cell depletion was comparably well tolerated and did not significantly affect the weight of infected guinea pigs or viral loads in their blood or tissue. However, significantly more viral genomes and transcripts were detected in the placenta and decidua of CD8^+^ T cell depleted dams post-infection. This study corroborates earlier findings made in nonhuman primates that maternal CD4^+^ T cells play a critical role in limiting the severity of primary CMV infection during pregnancy while also revealing that other innate and adaptive immune responses can compensate for an absent CD8^+^ T cell response in α-CD8-treated guinea pigs.

**Author Summary:** Congenital cytomegalovirus infection is a leading cause of adverse pregnancy outcomes and preventable disability in children. Using guinea pigs, a well-established small animal model of congenital infection and intrauterine development, this study tested how depleting T cells affects the course of primary cytomegalovirus infections. Severe illness and high rates of congenital infection were observed when helper CD4^+^ T cells were depleted. The depletion of killer CD8^+^ T cells did not affect the severity of disease or the rate of congenital infection but did increase the amount of virus that was detected in the placenta. A greater understanding how an immune response can prevent the infection of the placenta and developing offspring is needed to inform vaccine and therapeutic development. This study not only describes a new reagent that can be used to study the guinea pig immune system but also sheds new light in how adaptive immunity regulates congenital viral infection.

## Introduction

Human cytomegalovirus (HCMV), a ubiquitous beta-herpesvirus, is the leading cause of infectious disease-related birth defects and adverse pregnancy outcomes worldwide. Primary HCMV infection during pregnancy is ten times more likely to cause congenital infection than non-primary infections caused by either HCMV reactivation or reinfection [1], highlighting the key role of adaptive maternal immune responses in protecting against congenital cytomegalovirus infection (cCMV). How systemic maternal immunity and immune cells resident at the maternal-fetal interface (MFI) contribute to limiting congenital viral transmission remains critically understudied (reviewed in [2]). T cells play a critical role in maintaining immune homeostasis at the MFI, regulating the antiviral immune response, and in controlling HCMV reactivation or reinfection [3]. The combination of systemic and MFI-resident T cell responses may play key roles in preventing the transplacental transmission of HCMV [4, 5].

There are three distinct stages of T cell function during pregnancy and at the MFI: proinflammatory Th-1 T cell recruitment occurs during implantation and placentation, anti-inflammatory Th-2 T cells dominate between late first trimester and mid third trimester, and proinflammatory Th-1 T cells become more active during parturition [6, 7]. The transition from Th1 to Th2 immunity can enable some viruses, such as hepatitis E virus and influenza virus, to cause more severe infections during pregnancy [8–10]. Immune-suppressive CD4^+^ T regulatory cells (Tregs) play a critical role in maintaining immunologic tolerance to the fetus. In the mouse, immune adaptations that promote tolerance can make the placenta more susceptible to some bacterial infections [11, 12]. Abnormal Treg activity at the MFI, characterized by a shift in the abundance of inflammatory cytokine-releasing CD4^+^ T cells, is associated with preeclampsia, some placental infections, and spontaneous abortion [13–17]. Hyperactive CD8^+^ T cell responses at the MFI can also occur during preeclampsia, spontaneous abortion, and intrauterine growth restriction [18–23]. Maternal-fetal tolerance and the likelihood of successful pregnancies are negatively affected by the dysregulation of CD8^+^ T cell recruitment to and differentiation at the MFI; the systemic depletion of CD8^+^ T cells during pregnancy has similar effects [24–27].

Infection at the MFI can cause localized CD8^+^ T cell-mediated inflammation and, in severe cases, spontaneous abortion or fetal maldevelopment [28–30].

T cells play a critical role in antiviral immunity, and the effect of depleting T cells on primary cytomegalovirus (CMV) infections have been previously studied in mice and macaques. When mice were treated with CD4^+^ T cell-depleting monoclonal antibodies and infected with murine CMV (MCMV) there was no significant effect on the severity of MCMV disease, the kinetics of humoral immune responses were only slightly delayed, and CD4^+^ T cells were found to not directly control MCMV infection in multiple organs [31–35]. Others have reported that CD4^+^ T cells play key roles in controlling MCMV replication within the salivary glands, where CD8^+^ T cells and humoral responses are ineffective [33, 36–38]. Conversely, CD8^+^ T cells are primary effectors of anti-CMV immunity, as evidenced by the inflation of CMV-specific CD8^+^ memory T cells in response to primary infection and their role in controlling secondary lytic infection [39, 40]. While the depletion of CD8^+^ T cells did not affect the rate of MCMV clearance or cause severe disease, CD4^+^ T cells and NK cells gain costimulatory molecules and cytolytic activity in α-CD8 treated mice to compensate for the depletion of CD8^+^ T cells [33, 34, 41, 42].

The species-specificity of CMVs has presented a challenge to understanding the role of T cells in the pathogenesis of cCMV. Of the nonhuman viruses that are used to experimentally model HCMV infection *in vivo*, MCMV has been the most widely used but the virus does not cause placental or congenital infection in immunocompetent hosts [43–45]. Both rhesus CMV (RhCMV) and guinea pig CMV (GPCMV) cause placental and congenital infections and have been used for *in vivo* congenital CMV studies [46, 47]. Rhesus cytomegalovirus (RhCMV) infection during pregnancy is highly representative of congenital infection in humans; gestation and placentation in macaques and humans is highly similar, and RhCMV is more genetically similar to HCMV than GPCMV [48, 49]. However, use of the RhCMV model has been limited not only by the expense of experimental studies in nonhuman primates but also by the widespread endemicity of RhCMV in macaque colonies and limited available of CMV-seronegative animals of reproductive age [50–52]. Across two studies where seronegative macaques were infected with RhCMV during the late first/early second trimester, congenital RhCMV infection (defined as the detection of any viral DNA in amniotic fluid by qPCR) was reported in 47% (7 out of 15) cases [46, 53]. However, only one neonate from these experiments had congenital RhCMV-associated sequelae [46]. The depletion of maternal CD4^+^ T cells before RhCMV infection increased the rate of congenital infection to 100%, and fetuses were often either spontaneously aborted or died *in utero* [46, 53]. While RhCMV was detected in macaque placentas after infection by immunohistochemistry, an analysis of placentas from CD4^+^ T cell- depleted and immunocompetent dams did not find evidence that placental dysfunction had caused fetal death [46]. CD4^+^ T cell depletion delayed the production of RhCMV-specific humoral responses, but later studies found that preexisting humoral immunity in seropositive macaques or treatment with RhCMV-specific antibodies can protect against severe congenital infections after T cell depletion [53–55].

The guinea pig model of cCMV has been extensively used for preclinical CMV vaccine development since the virus was found to infect and transmit across the placenta [47, 56]. Guinea pigs have 65-day gestational periods, with fetal development and placental anatomy that is more similar to primates than murid rodents [57, 58]. Longitudinal studies of GPCMV infection during pregnancy have found that the virus infects the dam, placenta, and fetus sequentially and that necrotic lesions and hypoxic-like injuries develop in GPCMV-infected placentas [47, 59–61]. Maternal GPCMV infection late in gestation causes transcripts indicative of antiviral T cell responses to be upregulated in the placenta [62]. However, the limited availability of reagents for studying the guinea pig immune system has been a barrier to immunologic studies in the small animal [63, 64]. Monoclonal antibodies targeting guinea pig CD4 and CD8 that enable the targeted depletion of guinea pig T cells *in vivo* were recently developed [65–67]. In this study, we use these reagents to deplete CD4^+^ and CD8^+^ T cells during primary GPCMV infections in male and pregnant guinea pigs and evaluated the role of these T cell population in controlling infection and congenital transmission. Consistent with findings from the RhCMV model, our results show that CD4^+^ T cells are necessary for protection against severe GPCMV disease and the depletion of these cells leads to severe illness in adults and increased rates of congenital infection. The depletion of CD8^+^ T cells had no apparent effect on health or the rate of congenital infection despite causing elevated viral loads specifically at the MFI. Together, these experiments demonstrate the importance of T cell responses in preventing CMV from causing severe disease during pregnancy and establishing itself in or crossing the placenta.

## Results

### Subclass-switched antibodies targeting guinea pig CD8 deplete T cells *in vitro*

Monoclonal antibodies (mAbs) specific for guinea pig CD4 (H155, Rat IgG2a) and CD8 (B607 mouse IgG2a mAb), which were originally developed for immunostaining and flow cytometry applications, can deplete the expected T cell populations *in vitro* but are unable to deplete the lymphocytes for long durations *in vivo* [65–67]. Antibody effector functions are associated with IgG subclass, and a subclass-switched IgG2b variant of the α-CD4 antibody was developed and found to effectively deplete CD4^+^ T cells *in vivo* for extended periods [67–70].

The same subclass-switching approach was used to develop an IgG2b variant of the α-CD8 mAb. To confirm that the IgG2a and IgG2b variants of α-CD8 could deplete CD8^+^ T cells *in vitro*, guinea pig splenocytes were isolated and treated with 0.5 ug/mL of either mAb or purified rat IgG (rat IgG) and rabbit complement (none added or diluted to final concentrations of 1:6 or 1:12 complement). T cell abundance was measured by flow cytometry and the percentage of T cells that were depleted was calculated by comparing the abundance of CD8^+^ T cells in samples treated with rat IgG with samples treated with either α-CD8 IgG2 mAb (**Table 1**). Across two experiments, incubation with α-CD8 had no effect on T cells when complement was not added, but including either concentration of complement resulted in the depletion of >80% of CD8^+^ T cells in the sample. This experiment validated that the subclass-switched variant of α-CD8 retained the ability to deplete T cells *in vitro* by complement mediated lysis, as had been reported with the similarly modified α-CD4 [67].

**Table 1.** *In vitro* depletion of CD8+ T cells in guinea pig splenocytes following mAb treatment.

|  | % CD8<sup>+</sup> T-Cell Depletion±SD |  |  |
| --- | --- | --- | --- |
| **Complement** | **None** | **1:6** | **1:12** |
| **α-CD8 IgG2a** | -1.4±3.3 | 87.1±11.3 | 84.1±13.5 |
| **α-CD8 IgG2b** | 0.7±4.5 | 87.4±11.2 | 88.6±9.5 |

### The depletion of CD4^+^ but not CD8^+^ T cells increases the severity of primary GPCMV infections

We next assessed the effectiveness of α-CD8 mAb treatments on T cell depletion *in vivo* and evaluated the effect of depleting either CD4^+^ or CD8^+^ T cells on primary GPCMV infections in non-pregnant animals. Male strain 2 guinea pigs were infected with 1.0 × 10^6^ PFU of GPCMV by subcutaneous injection and treated with α-CD4, α-CD8, or rat IgG by intraperitoneal injection. The antibody treatments were repeated at 7- and 14-days post infection (DPI). Blood was collected at 0, 7, and 14 DPI to quantify GPCMV viremia by droplet digital PCR (ddPCR) and measure T cell abundance in PBMCs by flow cytometry (**Supplemental 1A**) [71]. Throughout this study, *in vivo* T cell depletion efficiencies were calculated as 1- (% circulating CD4^+^ or CD8^+^ cells at 7 DPI ÷ % circulating CD4^+^ or CD8^+^ cells at 0 DPI). All guinea pigs were euthanized and necropsied at 21 DPI; endpoint viral loads were measured in the blood and viscera by ddPCR and T cell abundance was measured by flow cytometry in isolated PBMCs and splenocytes.

In α-CD4 treated guinea pigs, an average of 99.4% of circulating CD4^+^ T cells were depleted at 7 DPI (**Fig. 1A**). The abundance of circulating CD4^+^ T cells increased at 14 and 21 DPI, either indicating that the potency of the mAb treatment was waning despite repeated injections and/or reflecting changes in T cell abundance that had occurred in response to the viral infection (**Fig. 1A**). The frequency of CD4^+^ T cells in splenocytes was significantly lower at 21 DPI after α-CD4 treatment, with a mean abundance of 9.1% in treated guinea pigs compared to 32.6% in rat IgG dosed controls (**Fig. 1B**). Primary GPCMV infections typically cause transient weight loss and peak viremia around 10 DPI; viremia generally resolves by 21 DPI after which point virus can be detected in visceral organs and salivary glands for weeks longer [63, 72]. Transient weight loss was observed in the rat IgG treated, GPCMV-infected group (**Fig. 1C**). The α-CD4 treated guinea pigs lost significantly more weight post-infection than the control group and did not recover their lost weight by the end of the experiment. Significantly elevated viremia was observed in the α-CD4 treated group as compared to the IgG treated group at 14 and 21 DPI (**Fig. 1D**). Where viremia resolved in the rat IgG treated controls by 21 DPI, viremia was observed in all CD4^+^ T cell depleted guinea pigs at the endpoint. Viral loads in the liver, lung, and spleens of the α-CD4 treated group were significantly higher than the rat IgG treated controls (**Fig. 1E**). The combination of prolonged and unresolved weight loss and elevated viral loads indicated that CD4^+^ T cell depletion increased the severity of primary GPCMV infections compared to control animals.

**Figure 1.**
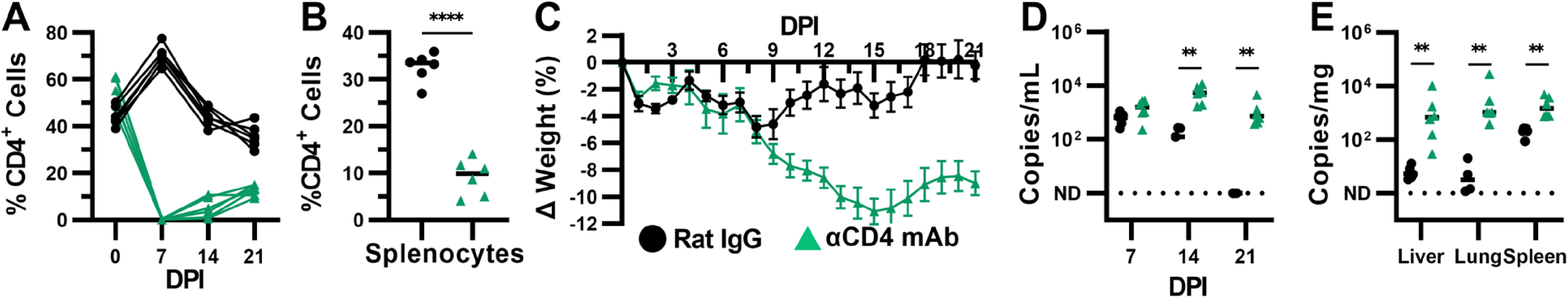
CD4^+^ T-cell depletion exacerbates primary GPCMV infections. Adult male guinea pigs were infected with 1×10^6^ PFU of GPCMV and treated with α-CD4 or rat IgG at 0, 7, and 14 DPI. The abundance of CD4^+^ cells among CD45^+^ PBMCs (**A**) and splenocytes (**B**) was determined by flow cytometry (**** p<0.0001, Mann-Whitney). (**C**) The mean change in guinea pig weight relative to 0 DPI over time. DNA was extracted from tissue and the abundance of viral genomes in whole blood (**D**) and tissues (**E**) was determined using a ddPCR assay specific to GPCMV *GP54* (** p<0.01, Mann-Whitney).

A second set of guinea pigs was used to study the effects of α-CD8 mAb and rat IgG treatments on GPCMV infection. As α-CD8 dosing for *in vivo* T cell depletion had not been optimized at the time of this experiment, we compared two treatments: 1.5 mg of IgG2b per dose or a 1:1 mixture of IgG2a and IgG2b (3 mg total/dose). No significant differences were observed between these two treatments and the results from the groups were combined for comparisons with the rat IgG-treated controls. As with α-CD4 treatment, the near-complete depletion of circulating CD8^+^ T cells was observed at 7 DPI, with an average depletion efficiency of 97.9%, although the abundance of these lymphocytes increased at 14 and 21 DPI despite the repeated antibody treatment (**Fig. 2A**). mAb treatment also significantly reduced the abundance of CD8^+^ T cells among splenocytes at 21 DPI (**Fig. 2B**). In contrast to the CD4^+^ T cell-depleted guinea pigs, α-CD8 and rat IgG treatments had similar effects on guinea pigs weight post-infection (**Fig. 2C**). CD8^+^ T cell depletion had no significant effect on GPCMV viremia or endpoint viral loads (**Fig. 2D, E**). Thus, while the depletion of CD4^+^ T cells heightened GPCMV viral loads and caused more severe disease in naïve guinea pigs, the depletion of CD8^+^ T cells was comparably well-tolerated and had no apparent effect on the acute phase of infection.

**Figure 2.**
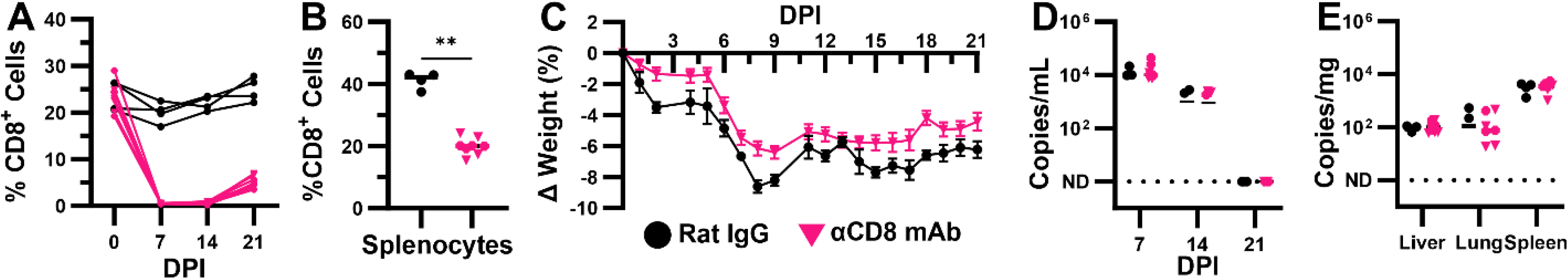
CD8^+^ T cell depletion has no apparent effect on primary GPCMV infections. Adult male guinea pigs were infected with 1×10^6^ PFU of GPCMV and treated with α-CD8 or rat IgG at 0, 7, and 14 DPI. α-CD8 IgG2b treatments are indicated with triangles and treatment with a mixture of α-CD8 IgG2a and IgG2b are indicated with crosses. The abundance of CD8^+^ cells among CD45^+^ cells in PBMCs (**A**) and splenocytes (**B**) was determined by flow cytometry (** p<0.01, Mann-Whitney). (**C**) The mean change in guinea pig weight relative to 0 DPI over time. DNA was extracted from tissue and the abundance of viral genomes in whole blood (**D**) and tissues (**E**) was determined using a ddPCR assay specific to GPCMV *GP54* (p>0.05, Mann- Whitney).

### CD4^+^ T cell depletion increases the severity of GPCMV infection in pregnant guinea pigs

Having confirmed that α-CD8 treatment depletes CD8^+^ T lymphocytes *in vivo* and compared effects of CD4^+^ and CD8^+^ depletions on GPCMV-infected male guinea pigs, we next assessed how T cell depletion affected mock- and GPCMV-infected pregnant guinea pigs. Strain 2 dams were bred with strain 13 boars during postpartum estrus to establish timed, semi-allogeneic pregnancies [73]. The dams (N=5/group) were infected with 1.0 × 10^6^ PFU of GPCMV or sham injected with PBS and treated with 1.5 mg of α-CD4, α-CD8, or rat IgG at 35 days gestation (dGA), which is past mid-gestation and during the fetal period of guinea pig development. At 7 DPI, the antibody injections were repeated, and the guinea pigs were bled so that circulating T cell abundance and viremia could be assessed.

We initially planned a 21 DPI endpoint to be consistent with the previously discussed experiments in male guinea pigs and because pathologic findings had been reported to be the most frequent in the GPCMV-infected placenta at this time [60]. However, three of five α-CD4- treated, GPCMV-infected dams succumbed to infection at either 13 or 14 DPI. This prompted us to end the experiment at 14 DPI so that the effect of T cell depletion on the fetuses and their placentas could be directly compared. We have found that guinea pigs bred during postpartum estrus typically gain 21±8.6% of their body weight between 35 and 49 dGA and all groups except the α-CD4, GPCMV-infected dams gained the expected amount of weight over the fourteen-day experiment (**Fig. 3A**) [62, 73]. Similar to what had been observed in male guinea pigs, the α-CD4, GPCMV-infected dams lost weight by the end of the experiment and were significantly smaller than dams from the other five groups.

**Figure 3.**
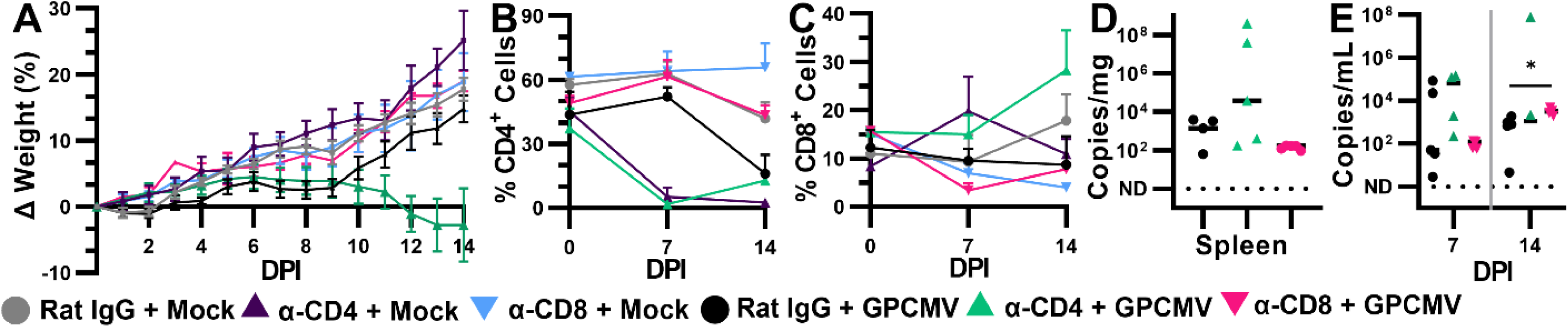
CD4^+^ T cell depletion and GPCMV infection causes severe disease during pregnancy. Time-mated guinea pigs were mock- or GPCMV-infected (1×10^6^ PFU) at 35 dGA and treated with α-CD4, α-CD8, or rat IgG at 0 and 7 DPI (N=5/group). (**A**) The mean change in guinea pig weight relative to 0 DPI over time. The abundance of CD4^+^ (**B**) and CD8^+^ (**C**) CD45^+^ PBMCs was measured by flow cytometry. DNA was extracted from tissue and whole blood, and the abundance of viral genomes in whole blood (**D**) and spleen (**E**) was determined using a ddPCR assay specific to GPCMV *GP54* (* p<0.05, Mann-Whitney).

The α-CD4 treatment was highly effective in pregnant guinea pigs, depleting an average of 99.4% of circulating CD4^+^ T cells at 7 DPI (**Fig. 3B**). α-CD8 treatment was somewhat less effective in pregnant guinea pigs relative to males, with mean circulating CD8^+^ T cell depletion efficiencies of 57.6% and 77.1% in mock- and GPCMV-infected dams at 7 DPI (**Fig. 3C**). CD4^+^ T cell-depleted dams had GPCMV viral loads that trended higher than the rat IgG-treated controls at 7 DPI and at necropsy (**Fig. 3D, E**). Blood could not be collected from the three dams that succumbed to infection, so it was unclear how α-CD4 treatment affected GPCMV viremia at 13-14 DPI. CD8^+^ T cell depletion had no significant effect on viremia or viral loads in the maternal spleen at 14 DPI. Taken together, CD4^+^ T cells depletion caused GPCMV infection to become more severe and potentially fatal in pregnant guinea pigs, as has been reported in RhCMV-infected macaques [46]. In contrast, CD8^+^ T cell depletion had no significant effects on maternal health or GPCMV viral loads.

### CD4^+^ and CD8^+^ T cell depletions have distinct effects on placental and congenital GPCMV infection

We next assessed the effects of T cell depletion and GPCMV infection on the MFI and fetus. The weights of fetuses and placentas were compared using mixed effects linear modeling to account for possible within-litter correlations caused by litter size, fetal sex, or the locations of conceptuses in the uterus. Placentas from α-CD4 treated, GPCMV-infected dams were significantly smaller than placentas from mock-infected dams treated with either rat IgG or α- CD4 (**Fig. 4A**). When viral loads were compared across the three groups of GPCMV-infected dams, α-CD8 treatment resulted in significantly elevated viral loads in the placenta and decidua compared to the rat IgG treated dams (**Fig. 4B**). While fetuses from the α-CD4 treated, GPCMV- infected litters trended smaller than the offspring of the other groups, no statistically significant differences in fetal mass were observed (**Fig. 4C**). CD4^+^ T cell depletion resulted in significantly higher viral loads in the fetal brain and liver than either rat IgG or α-CD8 treatment (**Fig. 4D**). Defining congenital infection as the detection of GPCMV in at least one tissue per fetus, 100% of the fetuses from the α-CD4 treated group were congenitally infected while congenital infection rates that averaged around 50% were observed in the control and α-CD8 treated groups (**Fig. 4E**).

**Figure 4.**
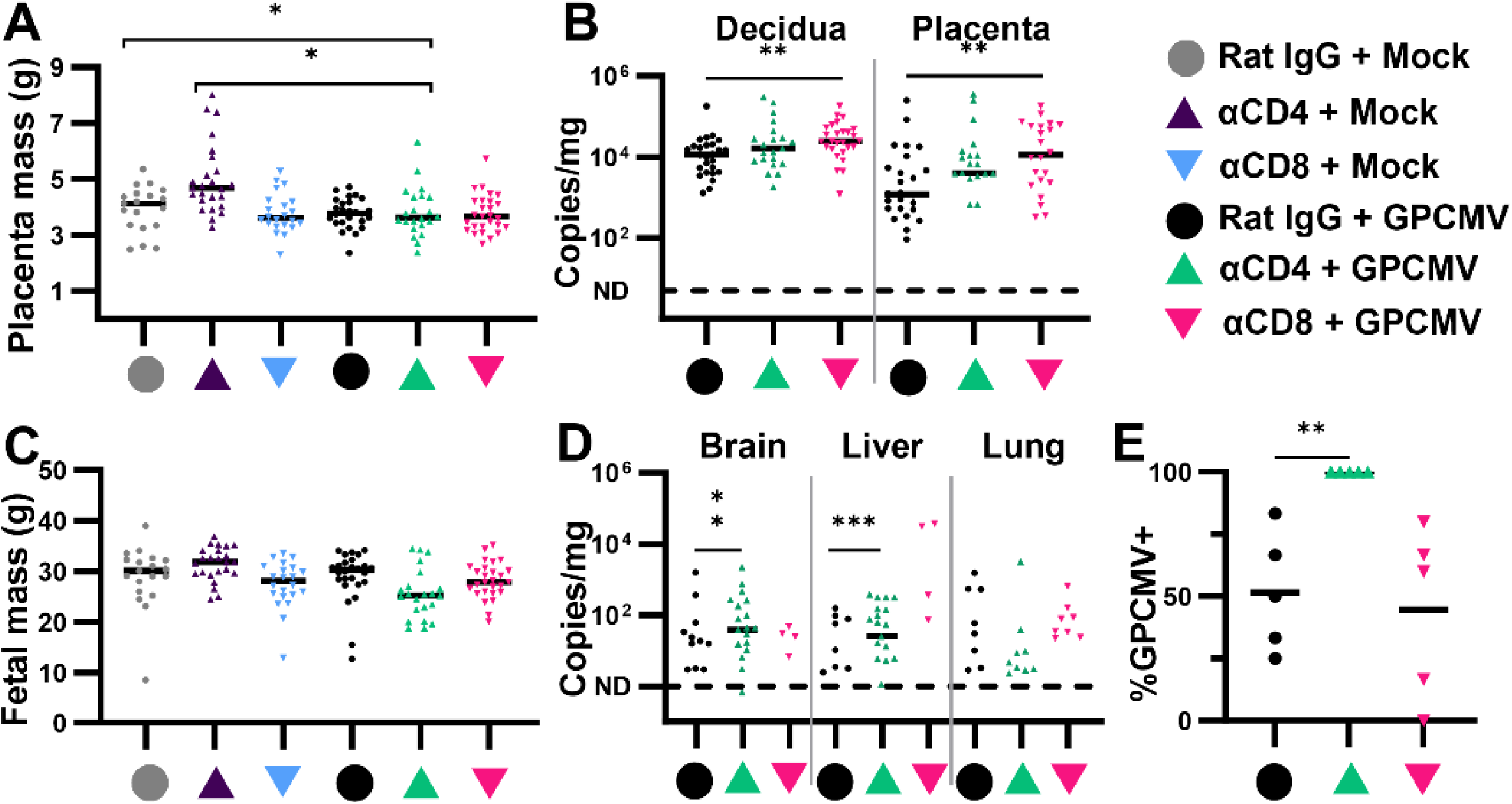
T cell depletion affects viral loads in the placenta and the rate of congenital infection. Time-mated guinea pigs were mock- or GPCMV-infected (1×10^6^ PFU) at 35 dGA and treated with α-CD4, α-CD8, or rat IgG at 0 and 7 DPI (N=5/group). Fetuses and placentas were collected after either maternal demise or necropsy at 14 DPI. The mass of placentas (**A**) and fetuses (**C**) were compared (* p<0.05, Tukey). DNA was extracted from tissue and the abundance of viral genomes at the MFI (**B**) or in the fetus (**D**) was determined using a ddPCR assay specific for GPCMV *GP54* (** p<0.01, Mann-Whitney). (**E**) The percentage of fetuses in each litter with congenital GPCMV infections, as defined as the detection of *GP54* in one or more tissues (**p<0.01, two-tailed T-test).

Finally, a histopathologic study was completed to assess the location of GPCMV-infected cells at the MFI and compare the effects of T cell depletion and/or infection on the placenta. Sections of hematoxylin and eosin (H&E) stained placentas were evaluated by a perinatal pathologist (**Supplemental Table 1**). The most severe pathologic findings occurred in two dams that had been treated with α-CD4 and had succumbed to GPCMV infection. All placentas recovered from one of these litters had globally infarcted within one day of maternal death. Five of nine placentas from the second litter had acute infarctions; fetal capillary death was apparent in all of the remaining viable placentas (**Fig. 5B**). The two CD4^+^ T cell-depleted, GPCMV infected litters that survived until 14 DPI had normal placentas and samples were not collected for pathology from the fifth dam in this group. Other placental abnormalities, including syncytiotrophoblast knotting and placental infarctions, were observed occasionally but were not clearly associated with infection or T cell depletion.

**Figure 5.**
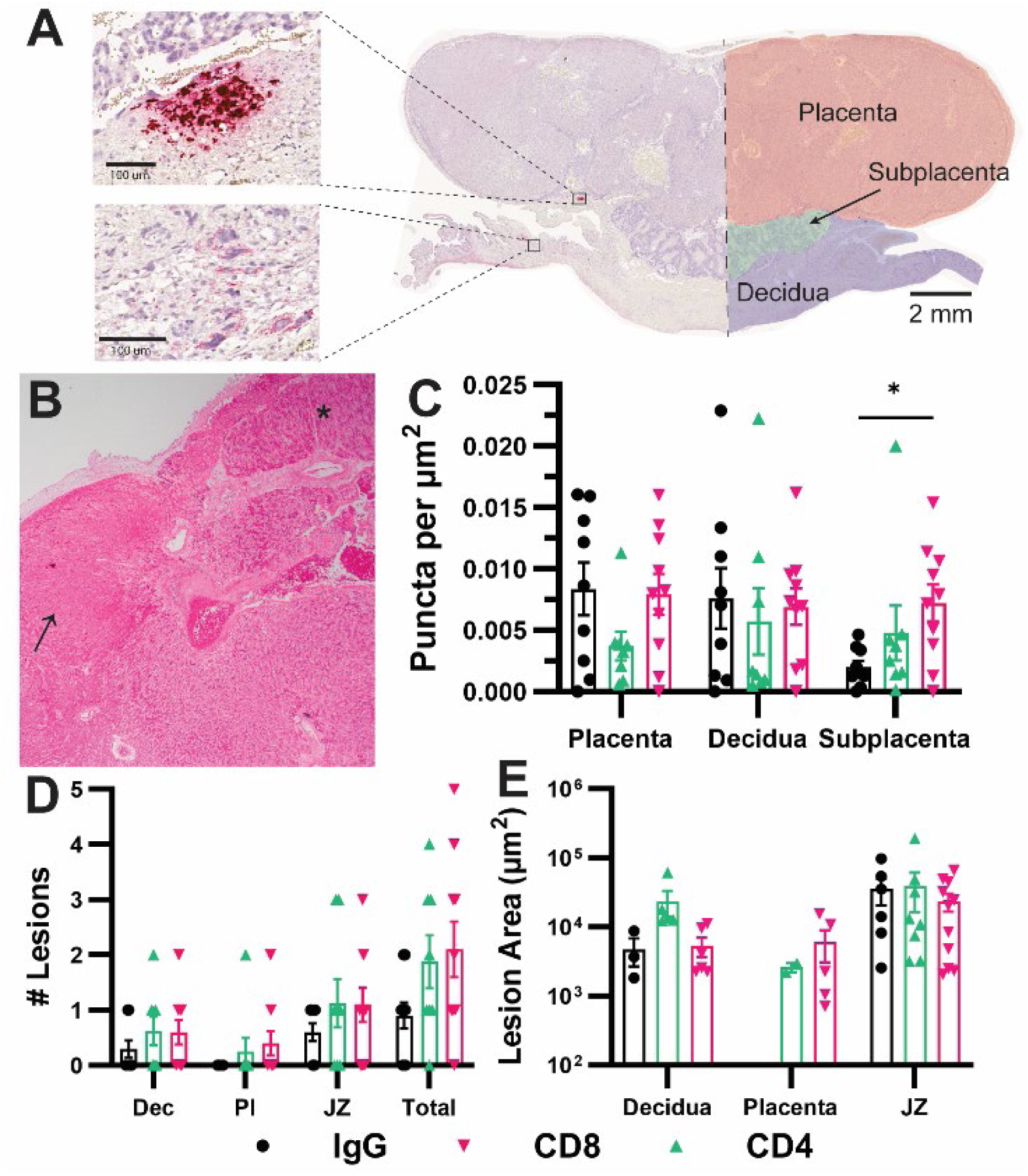
Histopathologic analysis reveals the effects of T cell depletion and infection on the placenta. Placentas were formalin-fixed and paraffin-embedded. Tissue sections were either H&E stained or stained by GPCMV-specific RNA-scope. (**A**) A representative image of *gp3*- stained placenta, showing tissue annotation for the automated detection of viral transcripts and representative areas of intense *gp3* staining (lesions) and puncta. (**B**) A representative micrograph of H&E-stained placenta from an α-CD4-treated, GPCMV-infected dam showing infarcted (arrow) and normal (asterisks) lobules. (**C**) The abundance of *gp3* puncta at the MFI normalized to area. (**D**) The number of intensely *gp3^+^* stained lesions detected in each placenta (Dec: decidua, Pl: placenta, JZ: junctional zone). (**E**) The area of each observed lesion was plotted and the extent of infection could be compared between the groups of placenta.

While ddPCR revealed that all the placentas and decidua recovered from GPCMV- infected dams were infected (**Fig. 4B**), this and other assays that quantify GPCMV abundance in tissue homogenate can overestimate placental infection by detecting virus or viral genomes that are present in maternal blood that was circulating through the placenta at the time of necropsy [60]. RNAscope specific to the viral transcript *gp3* was used to directly detect and compare the abundance of GPCMV-infected cells at the MFI [62, 71]. RNAscope is a highly sensitive method that can detect individual viral transcripts in cells, and we selected two placentas from each GPCMV-infected dam for analysis by RNAscope. Two patterns of *gp3* staining were observed (**Fig. 5A**). Intensely fast red-stained cells, where individual transcripts cannot be resolved, likely reflect productively infected cells, are rare, and are generally detected in the junctional zone or decidua [62]. Cells that contained fast red-stained punctate dots were much more frequent and could be found throughout many samples. To compare the abundance of viral transcripts across samples, an algorithm was used to detect red puncta at the MFI. Slide scans were manually segmented into the three major regions of the guinea pig MFI (the placenta, subplacenta, and decidua; **Fig. 5A**) and the abundance of puncta were normalized by area. In this analysis, GPCMV transcripts were more abundant in the placenta and decidua of rat IgG- and α-CD8- treated dams relative to animals treated with α-CD4, though this difference was not statistically significant (**Fig. 5C**). Significantly more *gp3* was detected in the subplacenta of α-CD8 treated- dams relative to the rat IgG treated controls.

A second, manual analysis compared the area and location of intensely Fast-Red stained “lesions”, which were defined as covering a minimum area of 500 µm^2^ and included at least three adjacent cells. For this analysis, we defined the junctional zone as tissue within 300 µm of the interface between the decidua and the main placenta or subplacenta. As we had previously reported at 21 DPI, most (56.4%, **Fig. 5D**) lesions were detected in the junctional zone [62]. The majority of the other lesions were found in the decidua, a few lesions were detected in the main placenta, and none were observed in the subplacenta. No significant differences in the size of lesions were observed between the groups (**Fig. 5E**). In conclusion, a combination of PCR- and *in situ* hybridization-based GPCMV detection suggests that CD8^+^ T cell depletion increases viral burden at the MFI, reflected as a higher abundance of viral genomes and transcripts, without affecting placental mass or the rate of congenital infection. CD4^+^ T cell depletion increased the rate of congenital infection, viral loads in fetuses, and caused severe placental abnormalities in cases where the infected dams succumbed to infection before necropsy.

## Discussion

The guinea pig is a valuable small animal model for understanding the developmental origins of disease and the pathogenesis of congenital CMV [47, 58]. Both humans and guinea pigs have hemomonochorial placentas that invade deeply into the decidua and both species develop to a similar extent during fetal life [57, 58, 74]. This study utilized recently developed mAb treatments to study the roles of CD4^+^ and CD8^+^ T cells in the immune response to primary CMV infection. We found that CD4^+^ T cells limit the severity of GPCMV infection in male and pregnant female guinea pigs and that the depletion of CD4^+^ T cells increases the rate of congenital infection, similar to findings made in the RhCMV model [46]. The depletion of CD8^+^ T cells was comparably well tolerated, though significantly higher viral loads were noted in the placenta and decidua of GPCMV-infected, α-CD8-treated dams.

The T cell response to CMV has been extensively studied in mice, and several studies have demonstrated that depleting or knocking out individual T cell populations does not result in severe MCMV disease as other immune cells can compensate for the lack of either CD4^+^ or CD8^+^ T cells [32, 35, 75]. Mice that lack normal CD4^+^ T cell responses have delayed antiviral responses but MCMV viremia eventually resolves as the virus establishes latency [76–78]. CD8^-^ CD4^-^ T cells can adopt some of the functions of CD4^+^ T cells in mice that are CD4^+^ T cell deficient, including mediating antibody class switching, supporting somatic hyper-mutation and affinity maturation of germinal center B cells, and assisting with a limited expansion of MCMV-specific CD8^+^ T cells [76–79]. The depletion of CD8^+^ T cells by monoclonal antibody treatment does not affect the rate of MCMV clearance, as humoral and natural killer cell responses can effectively control MCMV infections in the absence of a cytotoxic T cell response [33, 41, 42]. In pregnant macaques and guinea pigs, α-CD4 treatments cause primary CMV infection to be much more severe: many dams succumb to infection and the rate of congenital CMV transmission increases markedly [46, 53]. In a clinical setting, the rate of cCMV is significantly higher in HIV^+^ women [80–83]. While this observation reflects the key role of CD4^+^ T cells in preventing the vertical transmission of HCMV during pregnancy, only some of these studies found a correlation between the risk of cCMV infection and maternal CD4^+^ T cell counts.

This is the first study to examine how CMV infection during pregnancy is affected by depleting CD8^+^ T cells. α-CD8 treatment had no apparent effects on maternal health, maternal viral loads, or the rate of congenital infection. However, more virus was detected at the MFI in CD8^+^ T cell-depleted animals. Combining findings from this and a previous report, most GPCMV infected cells are detected at the junctional zone at 14 and 21 DPI when dams are infected after mid-gestation [62]. Why this region becomes sensitized to infection late in pregnancy and the cell types that are being infected remains unclear and merits further study.

Tissue-resident T cells in the maternal decidua may function as key immune barrier to vertical CMV transmission, but T cell responses to placental infection can also disrupt tolerance and compromise placental function [4, 5, 84]. For example, *Listeria monocytogenes* infection during mouse pregnancy can cause placental wastage and fetal demise, which can be prevented by therapeutically blocking T cell recruitment to the placenta or depleting maternal CD8^+^ T cells [12, 30, 85]. CMV infection can cause hypoxia-like injuries in the human and guinea pig placentas, though how the role of immune cell recruitment to the site of infection contributes to these injuries has yet to be fully elucidated [60, 86]. While we observed infarcted placentas in several α-CD4 treated, GPCMV-infected litters, we suspect that severe maternal illness is more likely to cause fetal harm than placental infection and dysfunction.

Ending our experiment at 14 DPI allowed us to compare viral loads and placental histopathology across the six experimental groups at a common time. However, 14 DPI may have been too early to detect adverse pregnancy outcomes or abnormal placental pathology caused by T cell depletion in uninfected guinea pigs or after α-CD8 treatment and GPCMV infection [60]. Follow up experiments will be necessary to determine how these treatments affect pregnancy outcomes, placental pathology, and to study T cell residence at the MFI in guinea pigs. This project was also affected by the effects of the COVID-19 pandemic on the global supply chain, and the sporadic availability of antibodies combined with a vendor’s loss of the α- guinea pig T Lymphocytes (Clone: CT5) hybridoma prevented us from using a consistent flow cytometry panel for all of our analyses. The scarcity of guinea pig-specific immunoreagents has been a long-term challenge for research using the species [64]. Given the importance of guinea pigs for infectious disease and perinatal research, a concerted effort should be made to increase the availability of existing guinea pig-specific antibodies and validate the functionality of other reagents for the small animal model [58].

## Materials and Methods

### Cells and Virus

GPCMV (SG13J2) was prepared as previously described [71]. Briefly, GPCMV strain 22122 (ATCC VR-682) was passaged 35 times in strain 2 guinea pigs between 1985 and 2010 [56, 87]. Salivary gland homogenate (SG13) was used to generate a seed stock by passaging the virus once on guinea pig lung fibroblasts (JH4, ATCC CCL-158). The seed stock was passaged once more on JH4 cells to prepare working stocks of virus that were used for this study. GPCMV stocks were titered on JH4 cells by plaque assay as previously described [88].

### Ethics Statement

All animal procedures were conducted in accordance with protocols approved by the Institutional Animal Care and Use Committee (IACUC) at the University of Minnesota, Minneapolis (Protocol ID: 2106-39180A). Experimental protocols and endpoints were developed in strict accordance with the National Institutes of Health Office of Laboratory Animal Welfare (Animal Welfare Assurance #A3456-01), Public Health Service Policy on Humane Care and Use of Laboratory Animals, and the United States Department of Agriculture Animal Welfare Act guidelines and regulations (USDA Registration # 41-R-0005) with the oversight and approval of the IACUC. Strain 2 and strain 13 guinea pigs were generously shared by Mark Schleiss and the U.S. Army Medical Research Institute of Infectious Diseases, respectively, and maintained in house as previously described since 2018 [62]. Guinea pigs were housed in a facility maintained by the University of Minnesota Research Animal Resources, who were accredited through the Association for Assessment and Accreditation of Laboratory Animal Care, International (AAALAC). All procedures were conducted by trained personnel under the supervision of veterinary staff.

### Monoclonal antibody production

Hybridomas that produce monoclonal antibodies specific for guinea pig CD4 and CD8 were developed by injecting rats or mice with guinea pig lymphocytes and fusing splenocytes with the mouse P3X63Ag8.653 myeloma line [65, 66, 89]. The H155 (rat α-CD4) and B607 (mouse α-CD8) hybridomas both express IgG2a. Spontaneous IgG2b subclass switch variants were developed using sib selection and ELISA as previously described [67, 70, 90]. CELLine 1000 Bioreactors (1000 ml scale, Wheaton) were used for monoclonal antibody production.

Hybridoma cells were suspended in Dulbecco’s Modified Eagle Medium (DMEM, ThermoFisher) supplemented with 10% IgG-depleted fetal bovine serum (FBS, Gibco, prepared by liquid chromatography on an ÄKTA pure [Cytiva] with a HiTrap Protein G HP column [Cytiva Product # 29048581]; IgG depletion was confirmed via lack of binding signal over baseline by biolayer interferometry on an Octet Red96e with Protein G biosensors [Sartorius]), 1% penicillin-streptomycin (Pen-Strep, Gibco), and 1% MEM non-essential amino acids (Gibco) and added to the cell compartment of the bioreactor. DMEM, supplemented as above except for FBS, was added to the medium compartment and the bioreactor was incubated at 37°C and 5% CO_2_. The bioreactor was initially seeded with 4.3 × 10^7^ cells in the cell compartment. Every seven days the DMEM in the medium compartment was replaced and the antibody-containing cell-side media was harvested. Cell viability was assessed by trypan blue staining, and 4.0 × 10^8^ cells in 15mL of cell-side media with a minimum viability of 30% was returned to the bioreactor. Monoclonal antibodies were purified from the harvested media via liquid chromatography on an ÄKTA pure with a HiTrap MabSelect PrismA protein A column (Cytiva Product # 17549851) (running buffer PBS, pH 7.4, elution buffer 0.1 M Na-Acetate, pH 3.5), then neutralized and buffer exchanged into Dulbecco’s Phosphate Buffer Saline (DPBS) [91]. Antibody concentrations were determined by absorbance at 280 nm using a Nanodrop spectrophotometer.

### Cell isolation and storage for flow cytometry

Leukocytes were harvested from male and female strain 2 guinea pigs. Peripheral blood mononuclear cells (PBMCs) were isolated from whole blood collected by toenail clip or cardiac puncture following euthanasia in K_2_EDTA blood tubes (McKesson). PBMCs were isolated by incubating 0.1 ml of whole blood with 0.9 ml of 1× red blood cell (RBC) lysis buffer (Thermo Fisher Scientific) for 10 to 15 min. The lysed blood was centrifuged at 300 × *g* and washed with DPBS three times. Isolated PBMCs were counted using Trypan Blue (Gibco) staining and a hemocytometer. The isolated PBMCs were either cultured in RPMI 1640 (ThermoFisher) supplemented with 10% FBS and 1% Pen-Strep (RPMI complete) overnight at 37 °C for immediate use or frozen in RPMI complete containing 10% DMSO (Millipore Sigma) for use within two years of their freeze date (1.0 × 10^7^ PBMCs/vial).

Splenocytes were isolated by harvesting spleens from euthanized guinea pigs and grinding spleens through a 100 µm cell strainer. The tissue was rinsed with 25 ml of RMPI complete and the cells were pelleted by centrifugation at 800 × g for 3 min. The cell pellet was resuspended in 10 ml of 1X RBC lysis buffer and incubated for 5 min at room temperature.

Splenocytes were pelleted by centrifugation at 500 × *g* for 5 min, the supernatant discarded, and cells resuspended in RPMI complete and counted using a Trypan Blue stain and a hemocytometer. Splenocytes were either cultured in RPMI complete overnight at 37°C for immediate use or frozen in RPMI complete containing 10% DMSO for use within two years of their freeze date (1.0 × 10^7^ splenocytes/vial).

### *In vitro* T cell depletion

The ability of monoclonal antibodies to deplete T cells by complement-mediated lysis was tested as previously described [67]. Guinea pig splenocytes were incubated with 0.5 µg of mAb in Hank’s Balanced Salt Solution (HBSS) and 5% FBS for 30 min at 4 °C. The splenocytes were centrifuged at 200 × *g* and resuspended in FBS-containing HBSS plus rabbit complement (BioRad) diluted to a final concentration of 1:6 or 1:12 or with no additional additions and incubated at 37 °C for 30 min. The cells were pelleted, washed once with DPBS, and cultured overnight at 37 °C in RPMI complete before T cell abundance was assessed by flow cytometry.

### Staining for flow cytometry

For flow cytometry analyses, leukocytes were cultured overnight to allow the potential re-expression of cell surface markers after monoclonal antibody exposure [67, 70]. Compensation controls were made from previously cryopreserved guinea pig PBMCs or splenocytes. Cells were centrifuged at 300 × g and resuspended in 1 ml of DPBS, and stained with Ghost Violet 510 (Tonbo Bioscience) viability dye for 30 min at 4°C in the dark. The cells were pelleted at 300 × g and resuspended in 0.1 ml of flow cytometry staining buffer (FACS buffer) (PBS pH 7.4 containing 0.5% bovine serum albumin (Thermo Fisher Scientific) and 0.1% sodium azide (Sigma Aldrich). 1 µl of Fc Block (α-mouse CD16/CD32, BD Biosciences) was added and the cells were incubated for 10 min at 4 °C in the dark. Cocktails of antibodies were prepared (α-PanT Lymphocytes:APC [Bio-Rad, MCA751APC], α-CD45:AF750 [Bio-Techne, NB100-65362AF750], α-CD4:PE [Bio-Rad, MCA749PE], α-CD8:FITC [Bio-Rad, MCA752F], α-CD14: PE-Cy7 [BioLegend 301814], and α-CD56:BV605 [BD 742659]), added to the Fc Block-treated cells, and incubated for 30 min at 4 °C in the dark. 1 ml of FACS buffer was added to the cells, the cells were centrifuged for 5 min at 300 × g, supernatant was aspirated, and the pellet resuspended in 2 ml of FACS buffer. Analysis was performed on a BD LSRFortessa H0081_X20 (configuration 093014) housed at and maintained by the University of Minnesota Flow Cytometry Core using the BD FACSDiva 8.0.1 (BD Biosciences). Flow cytometry data was analyzed using FlowJo v14.1 (Ashland, Oregon), T cell gating strategies are summarized in **Supplemental Figure 1**. Supply chain disruption caused by the COVID-19 pandemic and the loss of the α-guinea pig T lymphocyte hybridoma (Clone: CT5) by the manufacture resulted in an antibody panel that was not consistent throughout all experiments.

### GPCMV infection in T cell-depleted guinea pigs

Colonies of CMV-seronegative strain 2 and 13 guinea pigs were maintained in-house as previously described [62, 92]. ELISA was used to confirm the GPCMV serostatus of all animals before experimental infections were initiated [92]. To study the effect of T cell depletion on GPCMV infection in non-pregnant animals, four- to six-month-old strain 2 males were infected with 1X10^6^ PFU of GPCMV diluted in 0.5 ml in DPBS by subcutaneous injection into the scruff of the neck. At the same time, the guinea pigs were treated with 1.5 mg of IgG2b (α-CD4 or α- CD8), 1.5 mg of purified rat IgG (MD Biomedical), or 3 mg of a 1:1 mixture of IgG2a and IgG2b α-CD8 by intraperitoneal injection (N=4 or 6/group). These antibody treatments were repeated at 7 and 14 DPI. Blood and plasma were collected by toenail clip at 7 and 14 DPI, and the animals were euthanized at 21 DPI. Blood, plasma, spleens, liver, and lung were harvested at necropsy. Sample processing for flow cytometry and viral load quantification is described below.

To study the effect of T cell depletion in pregnant guinea pigs, male and female strain 2 guinea pigs were bred at 2 to 3 months of age. The strain 2 boar was replaced with a strain 13 male once fetuses were palpated so that timed pregnancies could be established by breeding the guinea pigs during postpartum estrus and produce semi allogenic pregnancies [73]. These timed pregnancies were confirmed by progesterone ELISA (DRG International); only animals with plasma progesterone concentrations exceeding 15 ng/ml by 21 days postpartum were utilized. At 35 days gestation, dams were injected with 1×10^6^ PFU of GPCMV or DPBS alone and treated with 1.5 mg of α-CD4, α-CD8 IgG2b, or rat IgG as described above (N=5/group). Blood and plasma were collected from the dams at 7 DPI, when the antibody treatments were also repeated. The guinea pigs were necropsied either after euthanasia at 14 DPI or after succumbing to GPCMV infection. Blood, plasma, and spleen were harvested from the dam. Fetal blood, brain, liver, lung and spleen were collected. Each placenta was divided, and half was immersion fixed with Shandon Formal-Fixx (Epredia) for 24 h and transferred to 70% ethanol until the tissue was paraffin-embedded. Portions of the remaining placenta and decidua were collected and frozen for viral load quantification.

### Viral load quantification and sexing by droplet digital PCR

DNA was extracted from whole blood or tissues using the DNeasy Blood & Tissue Kits (Qiagen). Viral genome abundance in samples was quantified by droplet digital PCR (ddPCR) using primers and probes specific to GPCMV *GP54* and the Bio-Rad QX200 system as previously described [71]. Hex-labeled primers and probes targeting guinea pig *Actb* (450 nM primer, 125 nM probe) and *Tspy2-like* (900 nM primer, 250 nM probe) were added to the reaction so that the sex of fetuses could be determined. With this amplitude multiplexing approach, female fetuses will have a single population of *Actb*^+^ droplets and male fetuses have additional, high-amplitude droplet populations that are either *Tspy2-like^+^* or *Actb*^+^ *Tspy2-like^+^*. After droplet generation, the ddPCR reactions were run using a C1000 Touch thermal cycler (Bio-Rad) and the following thermal conditions: 95 °C for 10 min; 40 cycles of 95 °C for 30 s and 56 °C for 60 s; 1 cycle of 98 °C for 10 min; hold at 4 °C. Droplet quantification results were analyzed using the QuantaSoft™ Analysis Pro software (Bio-Rad). GPCMV viral loads were calculated as the number of copies of viral genome per ml of blood or mg tissue.

### Placental histopathology

5-μm sections of formalin-fixed and paraffin-embedded placentas were prepared and mounted onto Superfrost Plus slides (ThermoFisher). Sections of each placenta were hematoxylin and eosin stained using standard methodology and analyzed by a perinatal pathologist for infection or T cell depletion-associated lesions. Up to two placentas from each litter were selected for GPCMV specific RNAscope [62]. Briefly, tissue sections were dried overnight and baked at 60 °C for 1 hr. Tissue was deparaffinized and pretreated using the recommended standard protocol for RNAscope® 2.5 Assays (ACD Document #322452). For target retrieval, the samples were incubated at 99 °C for 15 min, and the slides were treated with RNAscope Protease Plus for 30 min. Slides were stained using the RNAscope 2.5 HD Detection Reagent – RED (ACD Document # 322360-USM) using the V-CavHV-2-*gp3* RNAscope probe.

For RNAscope image analysis, stained slides were scanned using a Huron TissueScope LE at 20× magnification and visualized with QuPath (version 0.5.1) [93]. A script was developed for the automated detection of GPCMV transcripts. A pixel classifier was used to create a tissue thresholder and tissue boundaries were automatically annotated. Cells were identified based on nuclei segmentation from the hematoxylin optical density. The pixel size was 0.4 µm, the background radius setting was 8 µm, and the sigma setting was 1.5. The nucleus size parameters were set to 7 µm for the minimum and 400 µm for the maximum. The threshold level was 0.15, the maximum background was 2.0, and cell boundary expansion from the nucleus was 5.0 µm. Watershed post-processing was set to true. The subcellular spot detection module was used to detect fast red staining. fast red-stained spots were detected by spot minimal intensity and shape with an expected spot size of 0.5 µm^2^ and minimum and maximum spots sizes set to 0.5 µm^2^ and 2.0 µm^2^, respectively. Tissue was identified using the annotation setting; the minimum object size set to 2 × 10^7^ µm^2^ to select the tissue and avoid artifacts. Annotations were then split between the three regions (placenta, subplacenta, and decidua) using the wand tool, providing puncta counts and area for each annotated region.

For manual annotations, ≥500 µm^2^ areas that contained a minimum of three adjacent cells that were intensely fast red-stained (such that the algorithm could not count individual puncta) were defined as lesions. If the tissue had torn within or around the lesion the annotation boundary was drawn to exclude such space to provide an accurate determination of area [62]. Lesions that occurred within 300 µm of the main placenta or subplacenta meeting the decidua were defined as located within the junctional zone.

### Statistical Analyses

The weights of male guinea pigs were compared using a longitudinal linear model with a fixed effect for baseline (day 0). Maternal weights were compared using longitudinal mixed effects linear models, with fixed effect terms for day, treatment, day-by-treatment interaction, and baseline (day 0) weight, and random effect terms for intercept and slope by dam. Weight change trajectories were compared by treatment group by testing the day-by-treatment interaction term coefficients. The weights of fetuses and placentas were each examined using mixed effects linear models with fixed effect terms for treatment, sex, location within the uterus, and litter size, and a random effect term for the dam to account for within-litter correlations. Pairwise comparisons between pup mass and placental mass were adjusted for multiple comparisons (i.e. litter size, treatment) and compared using the Tukey method. Pairwise comparisons between viral loads and the abundance of viral transcripts (RNAscope puncta) were done using the Mann-Whitney rank comparison method. Analyses were conducted using R version 4.2.2 (R Foundation for Statistical Computing, Vienna, Austria). GraphPad Prism (v10.1.2) was used to graph data and for other statistical comparisons.

## Key Reagent Table

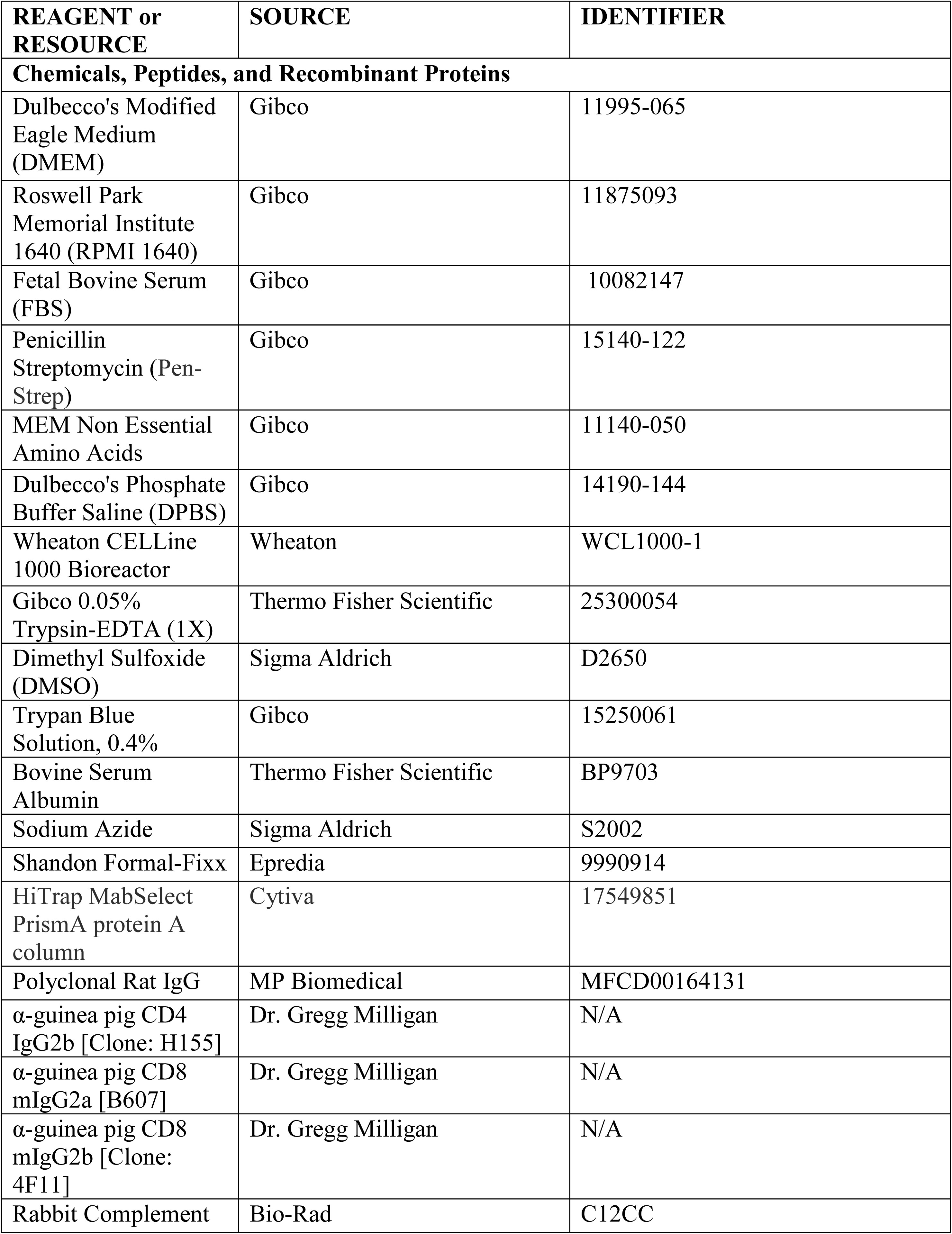
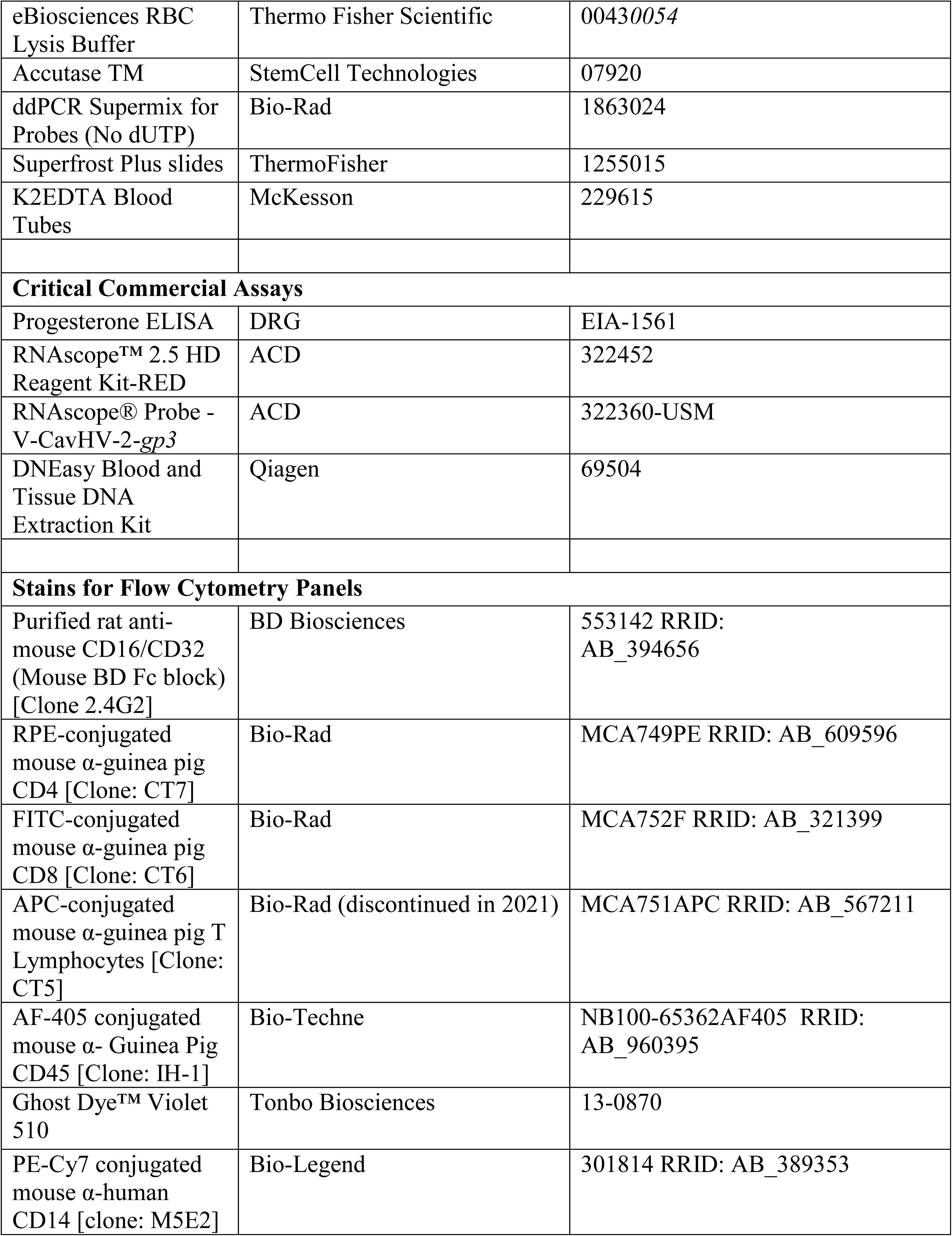
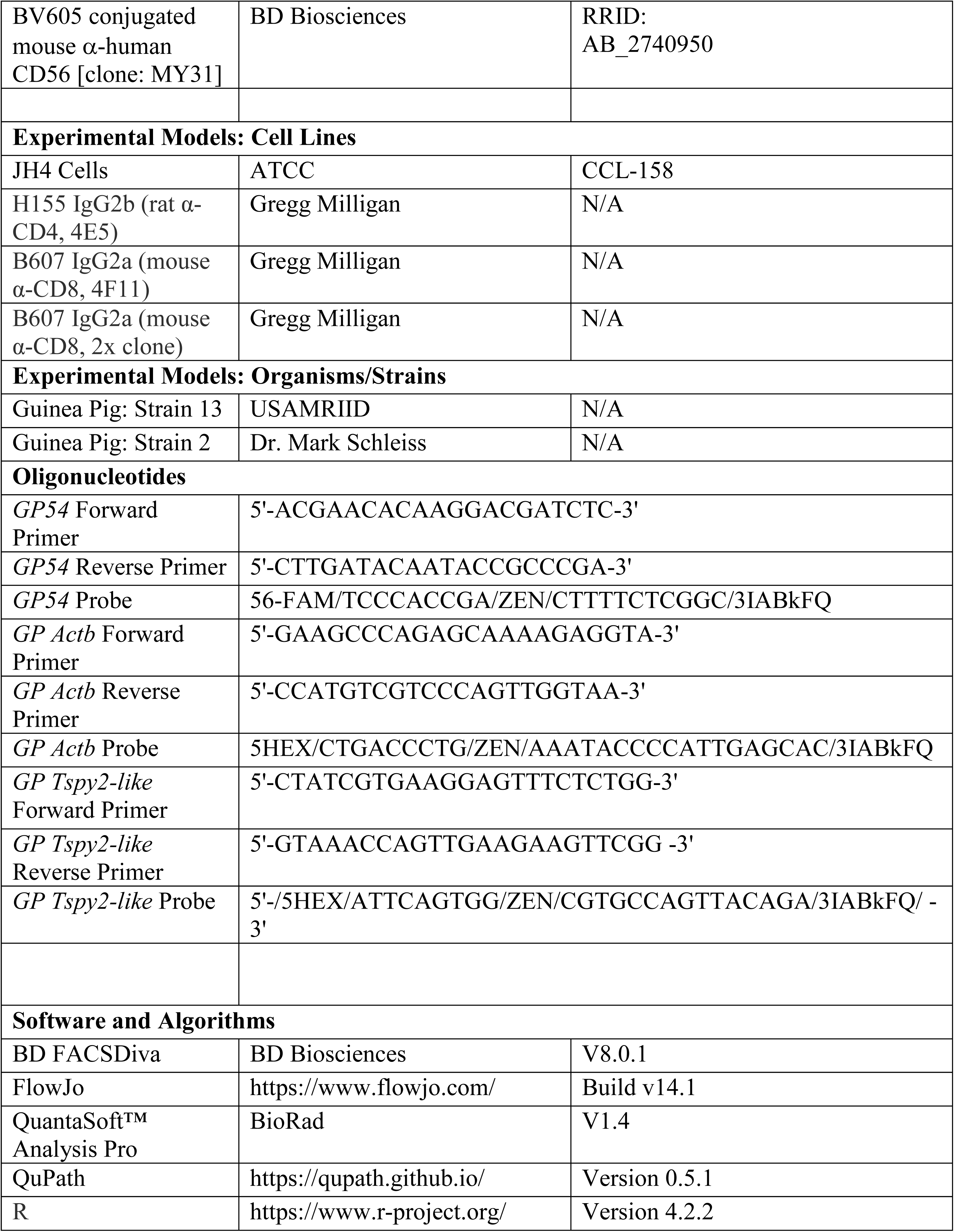

## Acknowledgements

This work was supported by the National Institutes of Health’s National Center for Advancing Translational Sciences, which provided funding for the Biorepository and Laboratory Services Team (Coleen Forster and Adam Lewis) and the Biostatistical Design and Analysis Center (Michael Evans). The University of Minnesota University Imaging Centers (SCR_020997, Mary E. Brown) and Flow Cytometry Resource supported microscopy and flow cytometry, respectively. The content of this article is solely the responsibility of the authors and does not necessarily represent the official views of the National Institutes of Health’s National Center for Advancing Translational Sciences.

---

**Supplemental Figure 1.**
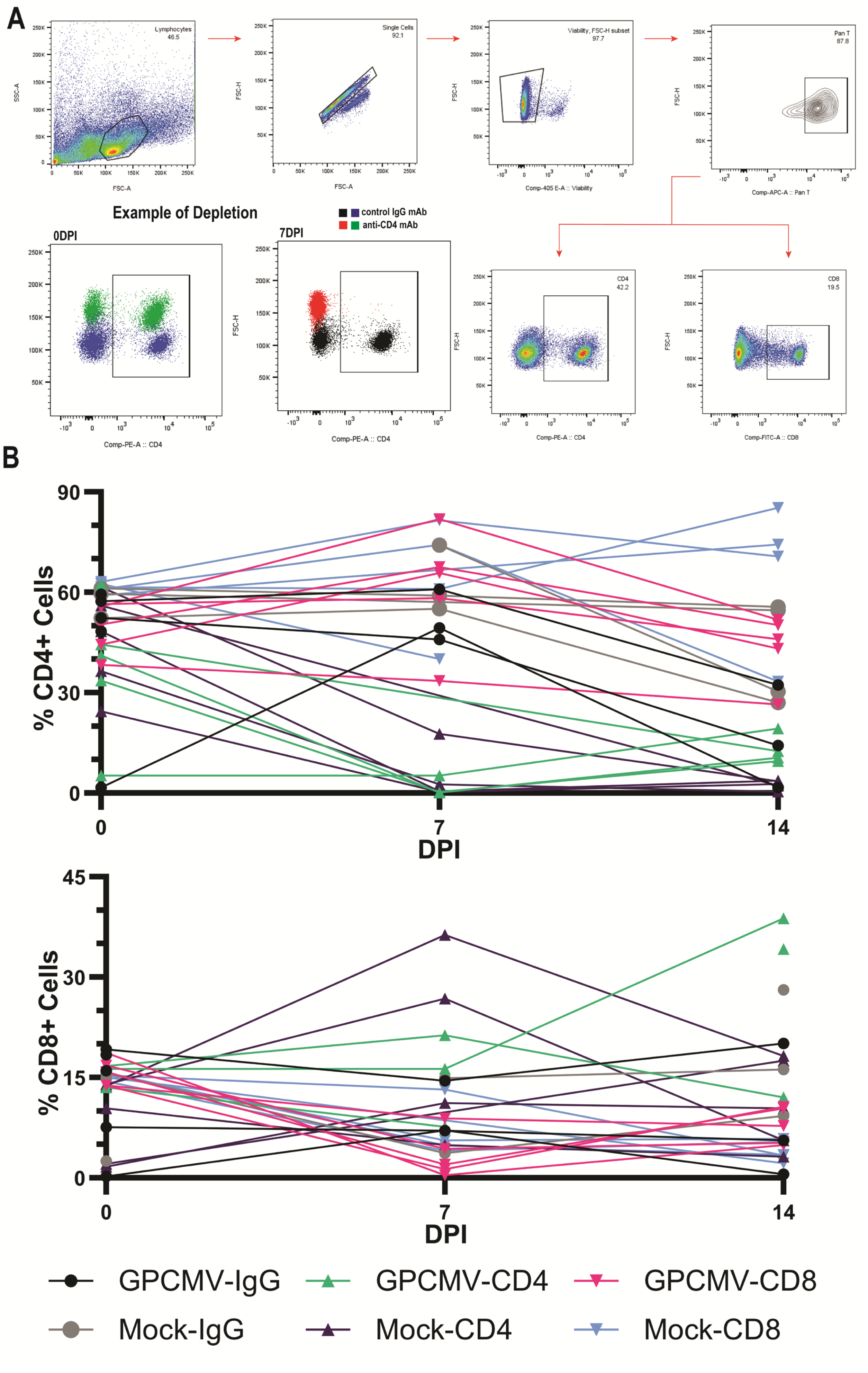
Flow cytometric analysis demonstrating the depletion of CD4^+^ or CD8^+^ T cells. Flow cytometry was used to quantify the abundance of CD4^+^ and CD8^+^ cells among CD45^+^ PBMCs or splenocytes. (**A**) Gating strategy used for this analysis and representative flow plots showing the effects of rat IgG and α-CD4 on PBMCs at 0 and 7 days post-treatment. (**B**) The abundance of circulating CD4^+^ or CD8^+^ cells in individual pregnant guinea pigs that were antibody treated and either mock- or GPCMV- infected.

**Supplemental Table 1. Data on individual pups and their placentas.**

